# Phenotypic and genomic comparison of human outbreak and cattle-associated Shiga toxin-producing *Escherichia coli* O157:H7

**DOI:** 10.1101/2022.09.30.510420

**Authors:** Nathan Peroutka-Bigus, Daniel W. Nielsen, Julian Trachsel, Kathy T. Mou, Vijay K. Sharma, Indira T. Kudva, Crystal L. Loving

## Abstract

*Escherichia coli* O157:H7 (O157)-adulterated food products, including beef and produce, are associated with disease outbreaks in humans. Although cattle feces are a source for the contamination, it is unclear if diverse O157 human-associated outbreak isolates expressing a specific virulence phenotype can colonize and shed in the feces of cattle at a quantitatively similar levels to non-outbreak isolates. It is also unclear if other phenotypes, such as biofilm, cell attachment, and toxin production, differentiate environmental O157 isolates from O157 isolates associated with human illness. Genomic profiling of O157 isolates acquired through routine surveillance can inform if the isolates encode virulence genes associated with human disease, but many genotype-phenotype relationships remain unclear for O157. Therefore, the objective of this study was to compare a diverse set of O157 isolates, with the intent of identifying potential genotypic differences that could inform phenotypes such as cattle colonization and fecal shedding, in *vitro* cell attachment, biofilm production, and Shiga toxin production. In addition, the relationship between phenotypes and potential for foodborne illness as it relates to genomic virulence traits was explored. No significant differences in cattle colonization and fecal shedding were detected for the tested isolates, despite broad genomic differences. In addition, the *in vitro* phenotypic differences noted in biofilm and cell attachment did not associate with one LSPA-6 lineage compared to another. Overall, no differences in cattle shedding were observed, yet variations in genotype and phenotype were identified indicating further work is warranted to better understand the relationship between O157 genome and virulence.

**Importance:** Foodborne illness has a major impact on the health and wellbeing of the global population, besides creating substantial financial hardships for industry. While many bacteria and viruses are implicated in foodborne illness, *Escherichia coli* serotype O157:H7 (O157) is a common food adulterant that can cause human disease and food recalls. Cattle feces are a significant source of food-adulterating O157. A greater understanding of O157 genetics and its relation to phenotype is needed to develop mitigation strategies to limit spread of O157 into the food chain. The goal of the research was to identify O157 genomic and phenotypic attributes of O157 associated with cattle colonization and fecal shedding along with other factors involved in environmental persistence and illness in humans. It was observed that variations in biofilm formation and *in vitro* cellular adherence did not associate with enhanced cattle colonization or fecal shedding, indicating that the processes involved in cattle colonization are complex and not well understood.

## Introduction

Foodborne illnesses in the United States are a significant burden to the welfare of the general populace, besides imposing considerable economic loss to industry groups. Shiga toxin-producing *Escherichia coli* (STEC) is a common cause of foodborne outbreaks. Illnesses associated with STEC are estimated to have an annual cost of 789 million dollars in the U.S, with O157 serotype responsible for 80% of these costs (1). Forty-three percent of STEC outbreaks during the years 2010–2017 were attributed to foodborne illness, with vegetable row crops (16%), beef (13%), dairy (10%), and fruit (4%) being the primary food sources (2). Foodborne illnesses associated with fresh produce have resulted in 83 multistate outbreaks in the U.S. from the years 2010-2017, resulting in 4,501 illnesses and 55 deaths (3). Of the 2010-2017 outbreaks associated with produce, pathogenic *E. coli* was responsible for 23 outbreaks, with O157 the dominant serotype behind 13 of the outbreaks and lettuce (i.e., romaine, iceberg, spinach, leafy greens) being the primary vehicle (3). Although, O157 is the most prevalent serotype associated with foodborne illness, other commonly implicated STEC serotypes include O26, O103, O111, O121, O145 and O45, which are all considered non-O157 STEC food adulterants by the USDA-Food Safety and Inspection Service (FSIS) (4). O157 still contributed to 23% of STEC illnesses during the years 2015-2017 (2, 5).

Cattle are a primary reservoir of O157, and O157 is often considered a commensal of cattle, as animals remain asymptomatic following STEC colonization (6, 7). O157 can be detected in various regions of the intestinal tract, but O157 predominately attaches to bovine epithelial cells at the recto-anal junction (RAJ) (8, 9) and is subsequently shed in feces. It poses a broad contamination risk, including beef products at slaughter and the environment. *E. coli* can establish itself as a member of many environments, including water, soil, and plants (10–12). Limiting O157 in cattle to minimize food contamination is important for preventing or reducing disease outbreaks in human (13).

Currently, O157 isolates are divided into three main phylogenetic lineages, (lineage I, lineage I/II, and lineage II) based on lineage-specific polymorphism assay 6 (LSPA-6) typing (14). Nucleotide polymorphisms in the lineage I and lineage I/II isolates are associated with pathogenesis in humans, such as a polymorphism in the *tir* gene at position 255, where a thymine (T) at this position is associated with clinical human isolates and an adenine (A) at this position is more associated with bovine isolates (15, 16). However, these broad generalizations are not all encompassing, as O157 isolates with *tir* 255 A alleles have been isolated from cases of human illness (15, 17). Current approaches in classifying O157 recovered from surveillance streams as potential isolates of concern for human illness includes the requirement for Shiga toxin genes (*stx2a*, *stx2c*, *stx1a*) along with the intimin gene *eae* (18). While consideration for toxin and intimin genes are important, these genes do not fully characterize different O157 isolates for the potential risk to human health or persistence in different environments. Thus, an improved understanding of the relationship between O157 genome and phenotypes expressed in various settings is important for utilizing data generated from various surveillance efforts.

We present various methods to evaluate O157 phenotypes with the intent to identify relationships between genome and phenotypes, particularly those associated with cattle colonization and human pathogenicity. Four O157 isolates from two different LSPA-6 lineages and sources were selected for analysis. The genomes were sequenced and their phylogenetic relationships to O157 isolates deposited in the NCBI Pathogen Detection database (https://www.ncbi.nlm.nih.gov/pathogens/) were assessed. Phenotypic assays consisting of cattle colonization and fecal shedding, *ex vivo* and *in vitro* cell attachment, biofilm formation, and Shiga toxin production were used to evaluate epidemiologically relevant phenotypes and their relationship to O157 pathogenic potential. In some cases, differences in phenotypes could be attributed to genomic differences, but much remains to be understood as attachment and biofilm phenotypes did not correlate to cattle colonization, nor did any of the O157 isolates differentially shed from cattle.

## Results

### Strain selection and comparison

*E. coli* O157 isolates were selected based upon LSPA-6 typing, isolation source, and association with human illness (Table 1). LSPA-6 lineage I isolates include EDL933 and TW14588, both of which are associated with foodborne outbreaks. LSPA-6 lineage I/II isolates included RM6067W and FRIK1989; RM6067W was associated with a foodborne outbreak, while FRIK1989 was isolated from dairy cattle feces and was not associated with human illness. Twenty-four-hour growth curves measured by changes in optical density in LB broth showed no major deviations between the four tested isolates; however, in low glucose DMEM, EDL933 achieved a lower plateau (Fig. S1).

**Table 1.** Shiga toxin-producing *Escherichia coli* O157 isolates used.

| Strain | LSPA-6<sup>1</sup> | Source | Outbreak Associated | Reference |
| --- | --- | --- | --- | --- |
| EDL933 | I | Ground beef (1982), USA | Yes | 80 |
| TW14588 | I | Lettuce (2006), USA | Yes | 78 |
| FRIK1989 | I/II | Dairy cattle feces (1991), USA | No | NADC<sup>2</sup> |
| RM6067W | I/II | Spinach (2006), USA | Yes | 77 |

<sup>1</sup>Lineage-specific polymorphism assay 6
<sup>2</sup>National Animal Disease Center, USDA

### Comparative genomics

To place our strains in the context of currently circulating O157 isolates, we compared genomic sequences from our strains to genomes in the NCBI Pathogen Detection Database https://www.ncbi.nlm.nih.gov/pathogens/. All isolates represented in the analysis possessed the major virulence genes *eae* and Shiga toxin (*stx*) genes and spanned LSPA-6 lineage I, I/II, and II. LSPA-6 types of modern isolates were largely congruent the with clades present in a core-genome based maximum likelihood phylogenetic tree (Fig. 1). The well-known foodborne outbreak isolates EDL933, TW14588, and RM6067W separated on the tree in clades with other clinical isolates from the NCBI Pathogen Detection database. The non-clinical isolate used in this study was isolated from a cow (FRIK1989), but related isolates were clinical isolates in lineage I/II. The currently circulating population of O157 strains is diverse and while the isolates represented in this study closely relate to isolates from human infections, they represent a small fraction of the broader phylogenetic diversity present.

**FIG 1.**
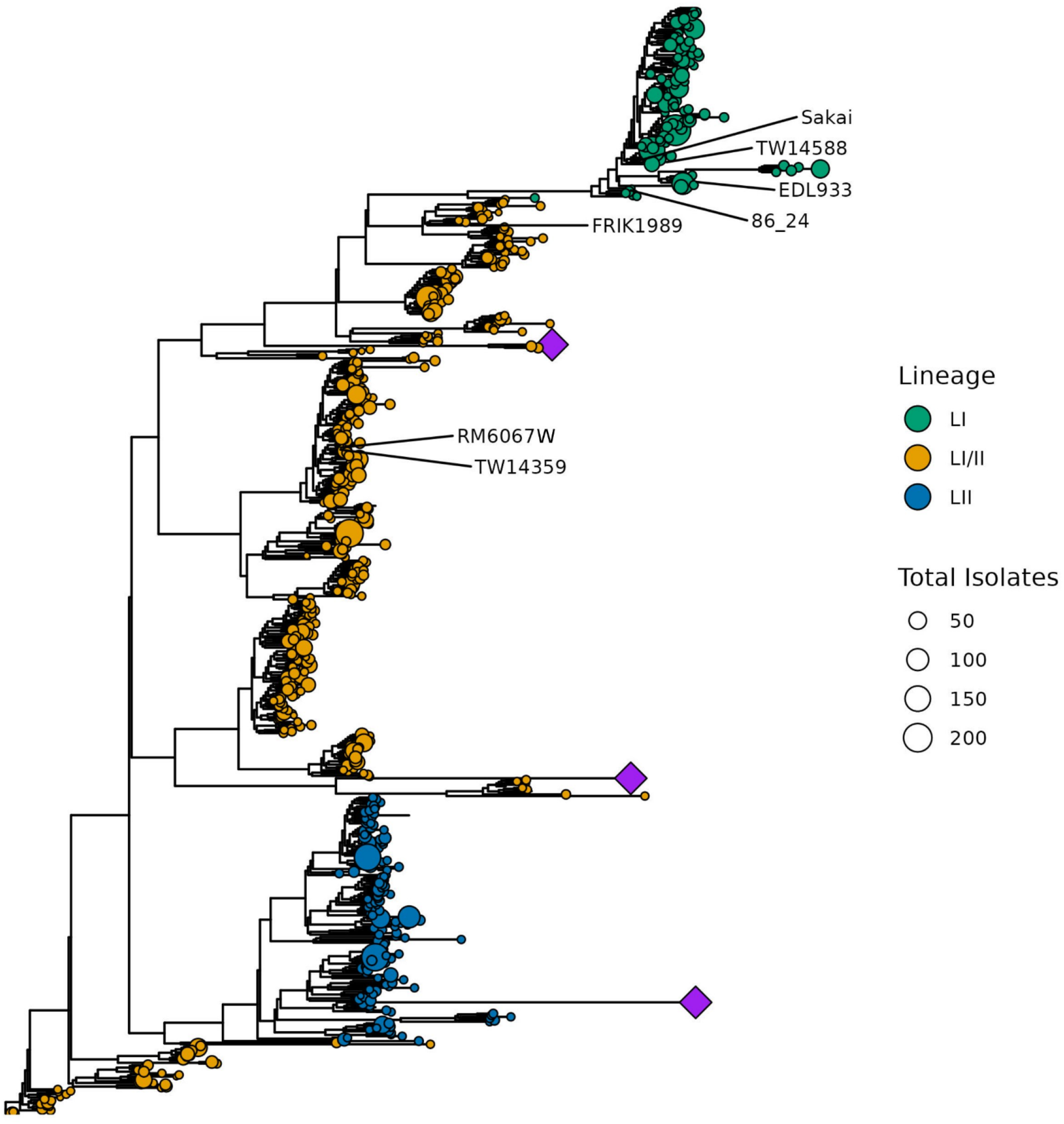
Phylogenetic relationship of *E. coli* O157 isolates used in this study. A maximum likelihood phylogenetic tree generated from a core genome alignment of 724 representative genomes (one per NCBI SNP cluster) and 7 reference genomes, include the 4 used in the current study. The circles representing the leaves of the tree are sized in proportion to the total number of genomes in that SNP cluster. The leaves are colored according to the LSPA-6 lineage typing scheme. Reference genome positions in the tree are indicated with text. Several clades exhibiting very long branch lengths were collapsed to improve the readability of the tree and are represented by purple diamonds.

Comparison of the four isolate genomes for the presence of known virulence genes revealed few differences, only the type of Shiga toxins genes and the Type 3 Secretion System (T3SS) effectors *espR4* and *ospG* (Fig. S2) were different across the four isolates. *Stx1a* was identified in EDL933 and FRIK1989, *stx2a* identified in all four isolates, and *stx2c* was detected in FRIK1989 and RM6067W. All four isolates possessed the T allele at position 255 in the translocated intimin receptor gene (*tir*), which is associated with pathogenesis in humans (15). The T3SS effector gene *espR4* was absent in FRIK1989 but detected in the other three isolates. RM6067W was the only isolate to possess the T3SS effector gene *ospG*.

The four O157 isolates could be differentiated using Polymorphic Amplified Typing Sequences (PATS) (19–21). Using PATS, genomic differences that conferred a distinct DNA fingerprint were identified as shown in Table S1.

### Fecal shedding and recto-anal colonization in cattle

Fecal shedding of the four O157 isolates from orally inoculated Jersey calves was analyzed using a trapezoid Area Under the Log Curve (AULC) for CFUs/g calculated from days 1, 2, 3, 4, 5, 7, 9, 11, and 14 post-inoculation. The PATS profile of isolates recovered matched that of the respective inoculum administered to the animals, thereby suggesting the lack of pre-existing O157. There was no significant difference in shedding among isolates over the 14-day period (*p* = 0.679, Fig. 2A). Time course of the average fecal shedding data for each of the isolates followed the same general pattern with a plateau in CFUs recovered at approximately days three to five post-inoculation and O157 recovered gradually dropped over the 14-day course of monitoring (Fig. S3A-D). It should be noted that by the end of the 14-day monitoring period not all calves had recoverable O157, with EDL933 having one of seven calves shedding, TW14588 having two of eight calves shedding, FRIK1989 having two of eight calves shedding, and RM6067W having four of seven calves shedding.

**FIG 2.**
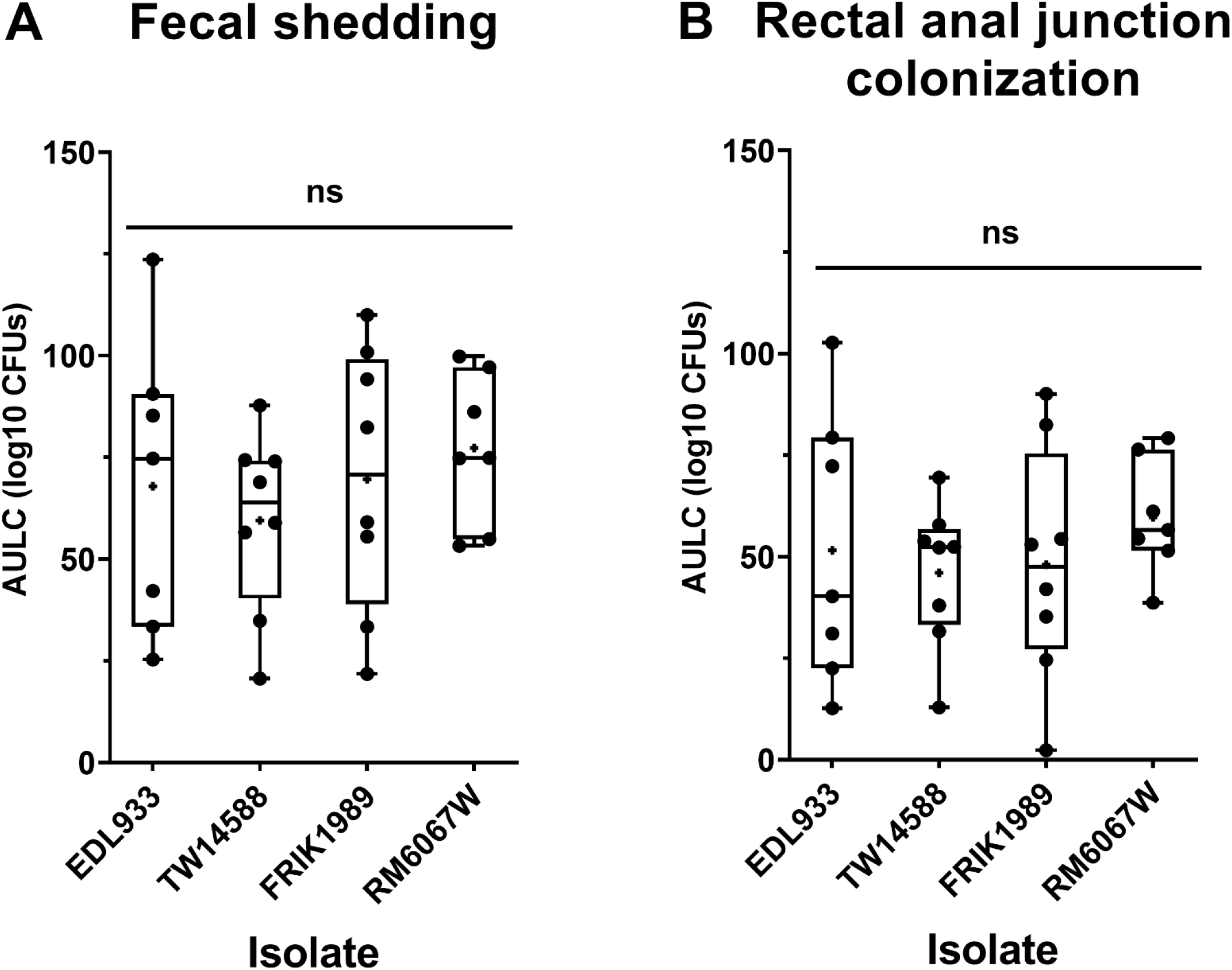
*E. coli* O157 fecal shedding and rectal anal junction mucosa colonization in Jersey calves. Jersey calves were orally inoculated with 6 x 10^9^ CFUs of indicated O157 isolates (EDL933, TW14588, FRIK1989, or RM6067W) and feces and recto-anal junction mucosa swabs (RAMS) were collected on days 1, 2, 3, 4, 5, 7, 9, 11, and 14 to enumerate O157 levels as indicated in materials and methods. Area Under the Log Curve (AULC) was performed (trapezoid Area Under the Curve model; RStudio version 2021.09.2). **(A)**. Cumulative fecal shedding and (B) cumulative RAMS colonization of respective STEC isolate. Each point represents an individual animal’s cumulative shedding/colonization in the indicated inoculation group. Statistical analysis was done by One-Way ANOVA with Tukey’s comparison of means; no significant (ns) difference in shedding or colonization, (*p* ≤ 0.05). Mean is indicated with a small “**+**”.

Differences in colonization at the RAJ was assessed through enumeration of O157 recovered from rectal-anal mucosal swabs (RAMS). Similar to fecal shedding, there were no significant differences in O157 in RAMS (*p* = 0.73, Fig 2B). Time course RAMS colonization trend for all isolates indicated average peak CFU counts from four to seven days post-inoculation, which gradually declined over the 14-day monitoring period (Fig. S3E-H). In a similar fashion to fecal counts, by the end of the 14-day monitoring period O157 was not recovered from the RAMS of all the calves in a challenge group, with EDL933 having three of seven calves still colonized, TW14588 having two of eight claves still colonized, FRIK1989 having three of eight calves still colonized, and RM6067W having six of seven calves still colonized.

### Assessment of O157 attachment to bovine epithelial cells

While the O157 isolates did not differentially colonize or shed from cattle, it’s unclear if shedding of these O157 isolates mirrors the *ex vivo*/*in vitro* phenotype, such as those resulting from attachment of these isolates to bovine cells. O157 isolates attached to *ex vivo* bovine RSE (RAJ squamous epithelial) cells (Fig 3), but the quantity and patterns of adherence were slightly different. As shown in Table 2, the O157 isolates demonstrated distinct attachment profiles compared to the control, non-pathogenic *E. coli* K12 isolate. All O157 isolates demonstrated some aggregation on RSE cells compared to *E. coli* K12, which attached in a diffuse manner. Of the four O157 isolates tested only FRIK1989 had a qualitatively strong attachment phenotype to bovine RSE cells (Table 2). The quantitative difference in RSE cell-attachment observed between all isolates tested was statistically significant (*p* <0.05) for isolates EDL933 (*p* = 0.0139), FRIK 1989 (*p* = 0.0076) and RM6067W (*p* = 0.0121), when compared to *E. coli* K12. All other comparisons did not yield any significant differences.

**FIG 3.**
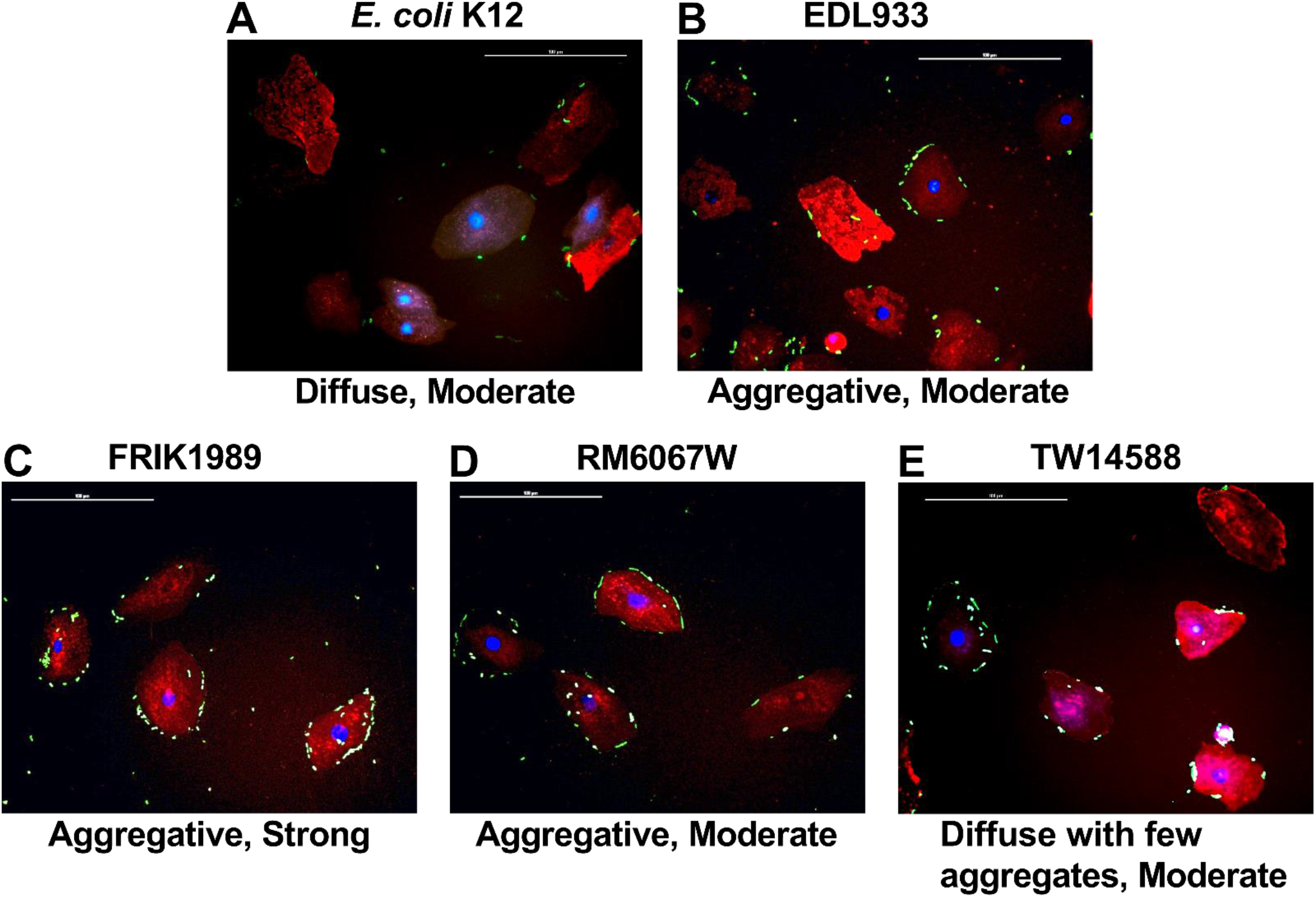
Adherence patterns of *E. coli* K12 and *E. coli* O157 isolates on RSE cells. *E. coli* isolates (K12, EDL933, TW14588, FRIK1989, and TW14588) were incubated with RSE cells at an MOI of 10:1 for 4 hours with agitation followed by assessment of cell attachment via immunofluorescence staining and co-localization. Representative immunofluorescent images with RSE cells and O157 are shown at 400x magnification. Adherence patterns on RSE cells were qualitatively recorded as diffuse or aggregative, with strong or moderate qualifiers as described in the materials and methods. *E. coli* were labeled with FITC (Green) conjugated antibodies. RSE cell cytokeratins were labeled with Alexa Flour 594 (Red) antibodies. RSE nuclei were stained with DAPI (Blue). Scale bar represents 100 µm. **Abbreviations**: Recto-anal Junction Squamous Epithelial (**RSE**), Multiplicity of Infection (**MOI**).

**Table 2.**
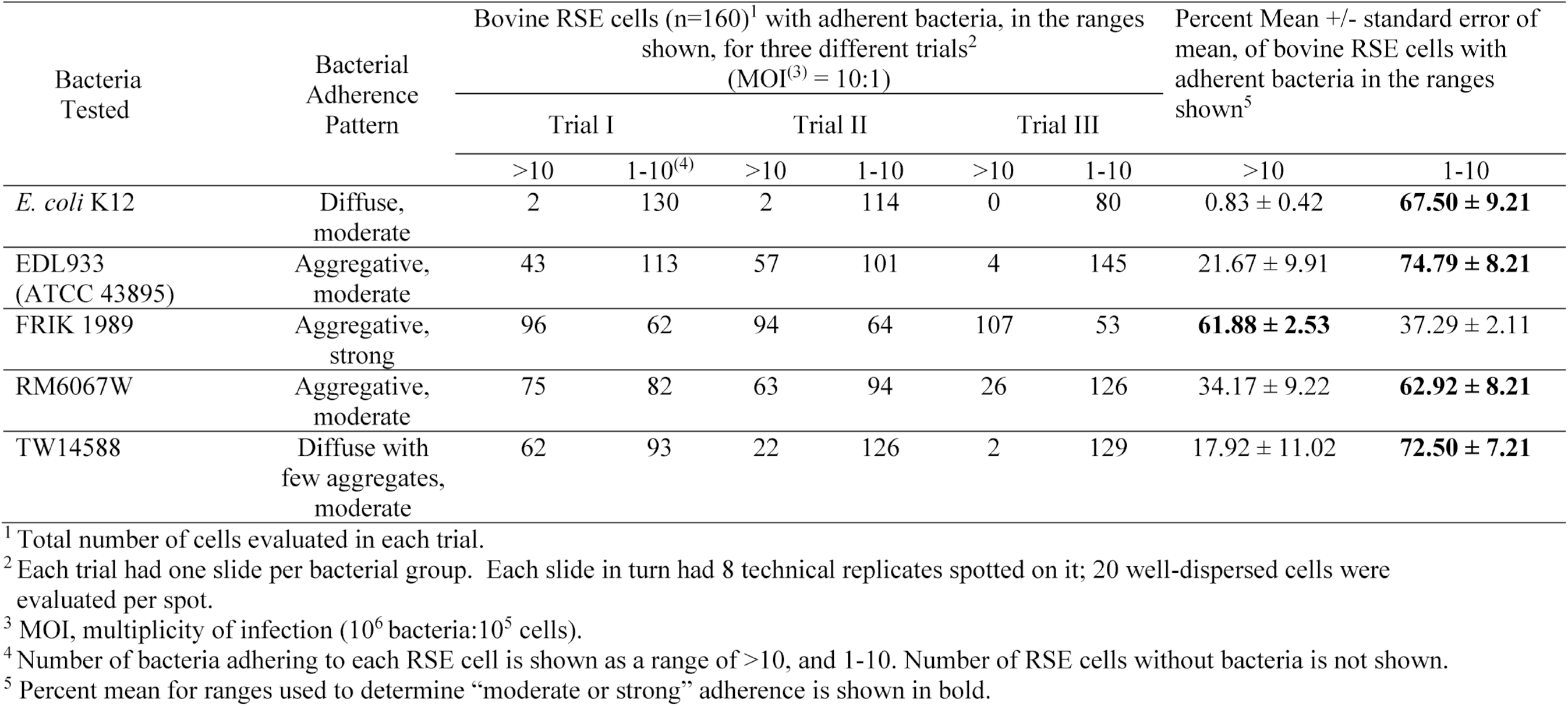
Quantitation of bovine recto-anal junction squamous epithelial (RSE) cells with adherent *E. coli* O157 bacteria.

Differences in attachment phenotypes were observed with the four O157 isolates when using a bovine intestinal epithelial cell (BIEC) line (Fig 4). EDL933 had the greatest attachment to BIECs, followed by FRIK1989, RM6067W, and TW14588. EDL933 attachment was significantly greater than that of the other tested isolates, with a 190-fold greater attachment to the BIECs than the non-pathogenic *E. coli* K12 (*p* < 0.0001), a 15-fold greater attachment than FRIK1989 (*p* < 0.0001), a 41-fold greater attachment than RM6067W (*p* < 0.0001), and vastly greater attachment than the other LSPA-6 lineage I isolate TW14588 (*p* < 0.0001) with a 252-fold increase in attachment. Of note, EDL933 formed microcolonies on the BIECs, while microcolony formation was not detected for the other O157 isolates (Fig. S4). FRIK1989 was the only other O157 isolate to display significantly greater attachment to the BIECs than *E. coli* K12, with a 11-fold greater attachment than *E. coli* K12 (*p* = 0.0066). Attachment of FRIK1989 compared to the other LSPA-6 lineage I/II isolate RM6067W was not significantly different. Attachment of TW14588 and RM6067W was not significantly different to that of *E. coli* K12.

**FIG 4.**
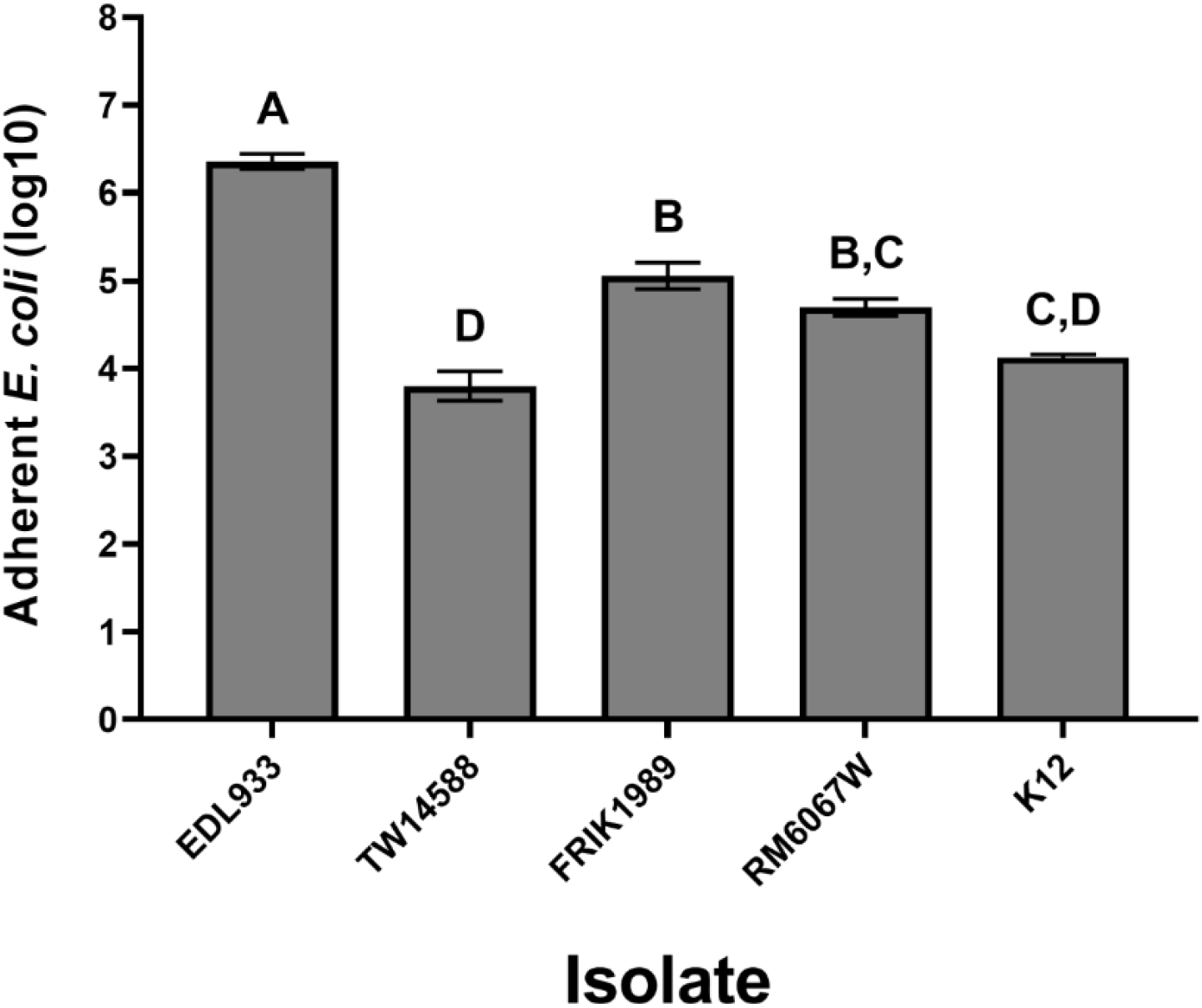
*E. coli* O157 *in vitro* adherence to bovine ileal epithelial cells. *E. coli* isolates (EDL933, TW14588, FRIK1989, RM6067W, or non-pathogenic K12) were incubated with bovine intestinal epithelial cells (BIEC, doi: 10.1007/s10616-018-0272-y) to assess epithelial cell attachment. The average number of bacteria initially added to the BIEC cultures across all experiments was 1.3 x 10^6^ CFU of each respective isolate. After 3h, non-adherent bacteria were removed via PBS washes and adherent bacteria were released from cells using 1.0% Triton X-100 with subsequent serial dilution and plating on LB agar for CFU enumeration. Values represent the Log10 mean ± the SEM of three independent experiments performed in triplicate wells for each experiment. A One-Way ANOVA with Tukey’s comparison of means was performed. Connecting letter report displayed above bars signify differences between *E. coli* isolate (*p* ≤ 0.05), with those bars with the same letter having no significant difference between them. K12 was included in only one of the three performed experiments.

### Biofilm production, curli expression, and Shiga toxin production

Biofilm formation is an important consideration for bacterial virulence and persistence in various environments. Different *in vitro* conditions can mimic various environments to inform associations between bacterial genotype and phenotype. Biofilm production was assessed for all four O157 isolates after propagation in both YESCA or BHI broth at both 26°C and 37°C utilizing either crystal violet staining or resazurin reduction (Fig. 5). YESCA broth was chosen as it is a low nutrient and low salt broth that promotes biofilm and curli production (22, 23), while BHI represents an enriched media. EDL933 produced robust biofilms under all test conditions. In YESCA broth FRIK1989 produced measurable, albeit small, amounts of biofilm at both temperatures, but failed to produce detectable biofilms in BHI at either temperature. TW14588 produced a small amount of biofilm in YESCA broth at 26°C, similar to FRIK1989. No biofilm production was detected under any of the assayed conditions for the RM6067W isolate.

**FIG 5.**
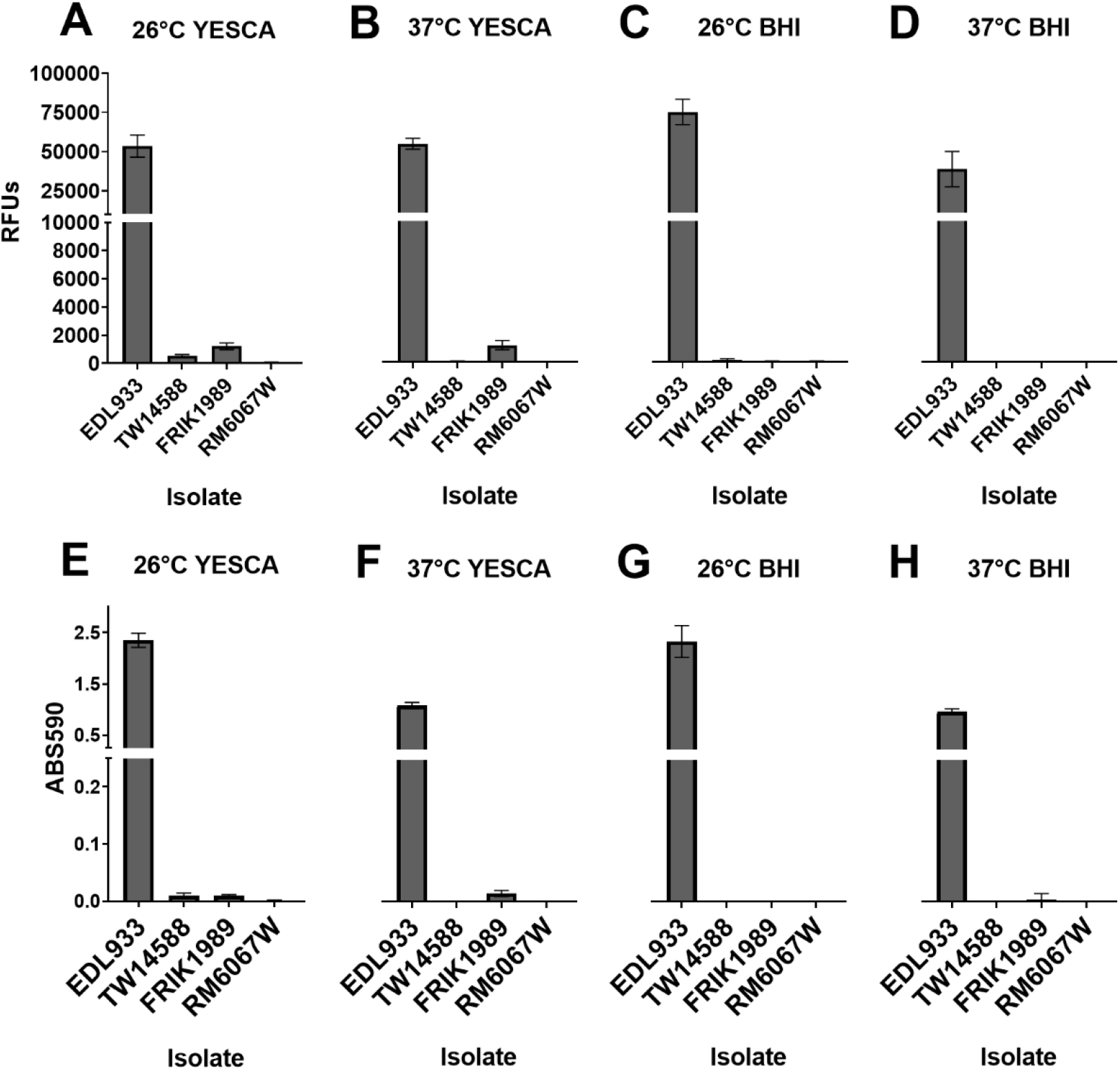
Biofilm production by *E. coli* O157 isolates under various conditions. *E. coli* isolates (indicated on x-axis) were cultured statically in 96-well microtiter plates for 5 days at either 26 °C or 37 °C in YESCA broth or BHI broth. Following removal of non-adherent bacteria, biofilm production was evaluated by either the viability reagent resazurin **(A-D)** or with crystal violet staining **(E-F)**. Resazurin indicates the viability of cells within the biofilm through the reduction of the non-fluorescent resazurin molecule to the fluorescent molecule resorufin via cellular NADP(H) dehydrogenase. Resazurin (0.03 mg/ml) was incubated with biofilms for 45 min and relative fluorescence units (RFUs) were measured using 530 excitation and 590 emissions; values shown are background subtracted. Crystal violet binds to both biofilm mass and bacteria indicating entire biofilm biomass. Crystal violet-stained biofilms were solubilized in 95% ethanol and absorbance was measured at 590nm (ABS590); a 1:5 dilution of the solubilized crystal violet was necessary as initial values obtained with the EDL933 isolate maxed out the detector. Values are background subtracted. Values represent the means ± the 95% CI of one experiment consisting of three independent cultures per isolate performed in triplicate. Y-axis for B-D are not shown, but are identical to A, and for F-H are identical to E **Abbreviations**: Relative fluorescence units (**RFUs**), Absorbance (**ABS**), Brain Heart Infusion (**BHI**).

Curli expression by O157, which can be detected with plating on Congo Red agar, is associated with biofilm production (24). EDL933 produced vibrant red colonies on Congo Red agar, while the three O157 isolates produced varying shades of pink, with RM6067W producing the lightest colored colonies (Fig. S5A-E). Biofilm production in *E. coli* O157 is in part regulated by the RNA polymerase sigma factor *rpoS*, and mutations in *rpoS* can result in biofilm production deficiencies (25–27). The *rpoS* gene for the four O157 isolates were translated *in silico*, and the resulting protein sequences aligned to the RpoS of *E. coli* K12 (Accession NP_417221); K12 was chosen as it’s a model biofilm producer and the role of its *rpoS* has been studied (28–30). The *rpoS* gene of RM6067W and FRIK1989 contained mutations that result in premature stop codons at position 135 for RM6067W and position 155 for FRIK1989, with the full-length protein at 331 amino acids (Fig. S5F).

Shiga toxin (Stx) production by O157 *E. coli* is a major virulence factor for human pathogenesis (31, 32). Stx production by each isolate was assessed using the Vero cell cytotoxicity assay (33, 34), as Vero cells are sensitive to Stx (Fig 6A). Stx cytotoxicity was inferred through the resulting loss in Vero cell viability as measured by resazurin for each isolate (33, 34). The greatest Vero cell cytotoxicity resulted from FRIK1989 supernatants with a 71% loss of Vero cell viability, which was significantly different to that of EDL933 (*p* = 0.0005), TW14588 (*p* = <0.0001), and RM6067W (*p* = <0.0001). EDL933 supernatants resulted in a 59% loss of Vero cell viability, which was significantly different compared to TW14588 (*p* = 0.0027) but not RM6067W (*p* = 0.4619). TW14588 and RM6067W produced similar levels of Vero cell cytotoxicity at 48% and 55%, respectively (*p* = 0.1253). No appreciable loss of Vero cell viability was detected with supernatants derived from *E. coli* K12, which does not encode *stx* genes (Fig. 6A). It is important to consider the susceptibility of Vero cells to the different Stx types, as different Stx types are associated with more severe disease in humans (34–36).

**FIG 6.**
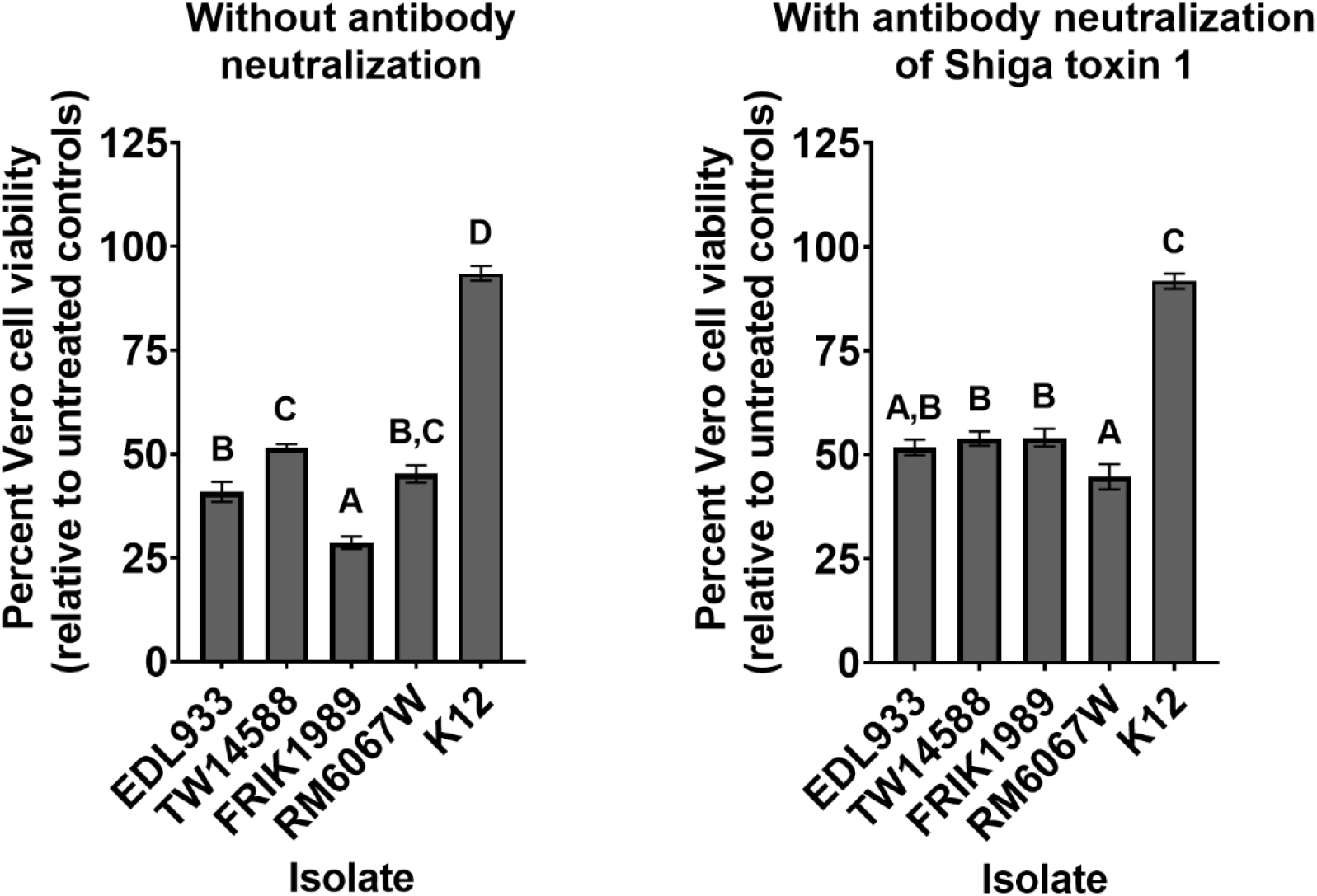
Vero cell cytotoxicity assay for Shiga toxin production by *E. coli* O157 isolates. Bacterial supernatants were collected from late log phase cultures (6 h) of indicated isolate grown statically at 37 °C in low glucose DMEM. Supernatants were added to confluent Vero cells for 48 h. Vero cell viability was determined using resazurin as described in materials and methods. Higher Shiga toxin (Stx) levels are inferred from decreased Vero cell viability, displayed as percent viable relative to non-treated Vero cell controls. **(A)**. Bar graph showing the percent viability of the Vero cells when treated with the 1:40 dilution of the *E. coli* supernatants with descriptive statistics. Values represent the means ± the SEM of two experiments performed in triplicate. Statistical analysis by One-Way ANOVA with Tukey’s comparison of means; connecting letter report displayed above bars signifies differences between O157 isolates (*p* ≤ 0.05). **(B)**. Bar graph showing the viability of Vero cells when treated with a 1:40 dilution of the bacterial supernatants incubated with Shiga like-toxin monoclonal antibody (13C4) for 2 hours at a 1:50 dilution to neutralize Stx1 from EDL933 and FRIK1989; isolates TW14588 and RM6067W lack *stx1*. Values represent the mean ± the SEM of two experiments performed in triplicate. Statistical analysis by One-Way ANOVA with Tukey’s comparison of means; connecting letter report displayed above bars signifies differences between *E. coli* isolates (*p* ≤ 0.05).

To evaluate the contribution of Shiga toxin 1 versus 2 to Vero cell cytotoxicity, an anti-Stx1 antibody was employed to neutralize Stx1 in supernatants prior to addition to Vero cells (Fig. 6B). When this approach was taken, RM6067W produced the greatest level of Vero cell cytotoxicity, with a 55% loss in Vero cell viability which was significantly different compared to that of TW14588 (*p* = 0.0437), and FRIK1989 (*p* = 0.0378), but not EDL933 (*p* = 0.1763). EDL933, TW14588, and FRIK1989 produced similar levels of Vero cell cytotoxicity at 48%, 46%, and 46% with Stx1 neutralized, respectively, which were not significantly different (*p* ≥ 0.5). The greatest change in Vero cell cytotoxicity with neutralization of Stx1 was for FRIK1989, the isolate that overall had the greatest level of cytotoxicity.

## Discussion

Understanding *E. coli* O157 genetics and its relationship to environmental survival and pathogenicity has been and continues to be important in protecting humans from foodborne illnesses (37–39). While considerable progress in inferring O157 pathogenesis from genomic data has occurred, the relationship between O157 genomic diversity and phenotypes expressed in various settings is still poorly understood (15, 40, 41). To better infer epidemiologically relevant attributes from diverse genotypes, there must be an input of phenotypic results to make genetic inferences. While the source of an isolate can provide some information, it does not provide sufficient information on the potential to cause disease and/or survive in the environment. For example, a collection of non-clinical O157 isolates demonstrated varying levels of Shiga toxin induced Vero cell cytotoxicity, with many isolates producing levels of cytotoxicity comparable to clinical and outbreak isolates (42). Lineage I, I/II, and II O157 isolates comprising both T and A *tir* alleles were recovered from cattle considered super-shedders (43). Varied biofilm and curli phenotypes from environmental O157 isolates are common, with some isolates producing both curli^-^ and curli^+^ colonies (44, 45). Surveillance of environmental isolates from cattle feces, soil, water, or produce is somewhat informative, but it must be acknowledged that being an environmental isolate does not preclude it from being pathogenic in humans. All four of the O157 isolates used here encoded the *tir* allele associated with human illness and also the *eae* and *stx2* genes, suggesting all four isolates could cause human illness if ingested (15, 46). While three of the isolates were associated with foodborne illness outbreaks (EDL933, TW14588, and RM6067W), the fourth isolate (FRIK1989) was isolated from cow feces but not associated with an outbreak (37, 47, 48). FRIK1989 belongs to LSPA-6 lineage I/II (often associated with human illness) and is closely related to clinical O157 isolates with sequence data in the NCBI Pathogen Detection database.

Understanding the relationship between O157 and its animal reservoir (e.g., cattle) is the first step in developing mitigation strategies to reduce O157 carriage in the animal and potential contamination of food products. There is great variation in reported shedding of O157 from experimentally inoculated calves across different studies (49, 50). The differences across studies could be due to different O157 isolates used as well as differences in experimental inoculation amounts, cattle breed, or pre-existing microbiota. In most studies, a single O157 isolate is used, which then limits comparisons of different isolates as it relates to shedding and/or colonization. The four O157 isolates used in the current study were separated both chronologically and by isolation source (37, 47, 48) and tested simultaneously in Jersey calves. Interestingly, no differences in colonization or shedding were detected. By day 14 post-inoculation, O157 was not recovered in feces or RAMS from many calves, with the exception of calves inoculated with RM6067W. O157 was still detected in six of the seven RM6067W inoculated calves on day 14. It is possible the tested O157 isolates may persist at or below the level of detection. It was observed that at times during the cattle challenge O157 were detected with RAMS, but was not detected in the feces, suggesting colonization will result in sporadic shedding and the absence of O157 in feces does not necessarily mean the absence of O157 in the gastrointestinal tract. Sporadic O157 shedding has been observed in longitudinal cattle herd studies (51). Overall, despite the genetic differences in the four O157 isolates, these did not result in differences in colonization or fecal shedding in cattle.

While animal studies are useful to understand the complex interactions between O157, commensals, and the animal host, they cannot be readily performed to assess O157 phenotypes. In addition, colonization and shedding are only a single phenotypic measure. Thus, attachment to bovine epithelial cells, including primary RSE cells (52, 53) or BIECs (54, 55) could serve as a surrogate to *in vivo* trials. O157 isolates that persist in cattle herds have strong attachment phenotypes (56). Thus, inferences may be made that O157 isolates with strong tissue culture attachment phenotypes may translate to greater colonization in cattle, but a relationship had not been clearly delineated. Tissue culture cell lines most frequently used to assess O157 attachment phenotypes are HEp-2, HeLa, or Caco-2 cells (57–61), which are all of human origin. In addition, HeLa and Hep-2 cells do not represent intestinal cells, and none are bovine cells. Caco-2 cells are derived from a patient with colorectal adenocarcinoma and are regularly used to study the intestinal epithelial barrier (62). Caco-2 cells may more closely approximate the intestinal environment, yet they are not representative of bovine. While ileum derived BIECs do represent a bovine intestinal cell, the ileum is not the preferred intestinal colonization site for O157, with main colonization observed at the recto-anal junction (9, 63). To better approximate recto-anal junction colonization, *ex vivo* bovine recto-anal junction squamous epithelial (RSE) cells were utilized to assess O157 attachment (52, 53).

EDL933 and FRIK1989 were the only isolates to adhere to BIECs in greater quantities than the non-pathogenic control *E. coli* K12, which lacks the ability to form intimin mediated pedestal formations on cells and has diminished attachment capabilities (64). FRIK1989, RM6067W, and EDL933 had greater quantitative attachment to RSE cells than non-pathogenic K12. While there are some common attachment phenotypes between these two cell lines, an association between O157 attachment to these cell lines and colonization in the cattle trial was not observed. The phenotypes derived from these *in vitro* assays could be useful in probing genetic differences among isolates, but further investigation is needed to address the association with the *in vivo* results. Of these four O157 isolates, it was those isolated from produce outbreaks (TW14588 and RM6067W) that had the weakest attachment phenotype; however, the screening of additional isolates will be needed to determine if this is a phenotype that can be universally associated with produce sourced O157 isolates.

Biofilm formation is problematic because it can facilitate transfer of O157 bacteria on food contact surfaces, promote their resistance to disinfectants, enhance their environmental survival, besides biofilms being hypothesized to play a role in cattle recto-anal junction colonization and shedding (65–68). Thus, biofilm formation and curli expression were assessed as additional phenotypes to explore differences across O157 isolates. EDL933 produced the greatest biofilm mass, showed highest curli expression, and strong cellular attachment phenotype. A link between biofilm production and cellular attachment has been proposed (69). EDL933 also forms microcolonies, which may be related to biofilm production (70). FRIK1989 displayed a strong attachment phenotype yet was a low biofilm producer and low curli producer, indicating that additional elements are involved in cellular attachment versus biofilm formation. Kudva et al. described curli variants with differential attachment patterns to RSE versus HEp-2 cells (23), and results indicated O157 proteins involved in attachment to bovine cells differed from those involved in attachment to human cells. Biofilm production may be more relevant in environmental persistence (68), which would be advantageous to the organism in disseminating itself along production chains. Even in the absence of biofilm forming capabilities in O157 isolates, these biofilm deficient isolates will integrate themselves in mixed-species biofilms and confer a survival advantage (71). Interestingly, EDL933 curli deficient mutants were observed to have poorer persistence on spinach leaves than curli expressing EDL933 (72), and yet we observed the produce-associated isolates (TW14588 and RM6067W) to be poor curli producers. Others have observed that curli expression doesn’t impact internalization into spinach roots using mutants of the 86-24 O157 isolate (73). This suggests that biofilm and curli expression likely contribute to produce colonization but is dependent on the specific interactions between bacteria and the plant environment. The absence of a strict association between biofilm formation and virulence, as seen here with these O157 isolates, is also observed with the pathogenic bacteria *Listeria monocytogenes* (74).

Shiga toxin production is an important consideration for O157 virulence in humans (46). While *stx*-encoding genes suggests the potential of O157 to cause disease in humans, the toxin gene type and expression levels of encoded toxin, which to some degree is linked to the lytic nature of the Shiga toxin encoding phage, are important considerations as it relates to human disease (75). Stx2 is associated with more severe clinical outcome (34–36). The FDA’s relative risk index for O157 isolates ranks isolates in order of potential severity in human illness by *stx* genes with *stx2a* > *stx2c* > *stx2a* + *stx1a* > *stx1a* (18). The four O157 isolates studied have different *stx* gene profiles (Fig. S2). RM6067W had the most hemolytic uremic syndrome (HUS) cases associated with its outbreak (76–78) and possessed both *stx2a* and *stx2c*. FRIK1989 also possessed *stx2a* and *stx2c*, and it was observed that it’s *stx1a* expression was a major contributor to the Vero cell cytotoxicity. The Vero cell assay for Shiga toxin expression seems like a suitable model to infer potential adverse clinical outcome of O157 isolates when Stx1a is neutralized. However, it is generally assumed that O157 isolates that cause Vero cell cytotoxicity have the potential to cause illness in humans. In the absence of *Stx* gene expression data, only assessing for the presence/absence of various *stx* genes may not be sufficient to inform virulence potential of various isolates.

Overall, diverse phenotypes within *E. coli* O157 are appreciated, yet a clear phenotypic relationship to cattle shedding and human disease have not been fully identified. Phenotypic heterogeneity is attributed to improved fitness in virulent bacterial populations (79), and could help explain the variation of observed phenotypes between these *E. coli* O157 isolates. The phenotypic and genetic determinants involved in cattle colonization are complex and warrant further study for development of intervention strategies. Shiga toxin production, not just gene content, remains an important phenotype to infer clinical virulence of O157. However, a defined threshold of Stx production that would result in clinical illness is unclear. One limitation of this study is the small number of O157 isolates used. Evaluating isolates of known persistent cattle colonization and/or additional clinical isolates would enable better inferences into tested phenotypes and how they relate to cattle colonization and fecal shedding along with environmental survival and virulence in humans. Inter-animal variation in colonization and fecal shedding among the cattle potentially confounded the results, indicating future studies should include a greater challenge cohort size to account for this variability. While no significant differences in cattle colonization and fecal shedding were observed between the four O157 isolates, TW14588 cattle colonization and fecal shedding counts were on average lower than the other O157 isolates and at the same time TW14588 attachment counts to BIEC and RSE cell lines were lower than the other tested O157 isolates. O157 attachment to BIEC and RSE cells could be useful in predicting cattle colonization and fecal shedding if less inter-animal variation is observed or a greater number of animals are used. Testing additional phenotypic profiles, such as carbon source utilization or acid resistance, may fill in the gaps presented here and help in finding additional phenotypes that can correlate to animal carriage or environmental survival.

## Materials and Methods

### Isolate selection and cultivation

Four O157:H7 isolates (EDL933, TW14588, FRIK1989, and RM6067W) were selected for analysis in this study based on their LSPA-6 lineage and isolation source (Table 1) (77, 78, 80). *E. coli* K12 was used as an avirulent control in *in vitro* experiments. *E. coli* isolates EDL933 (ATCC 43895) and K12 (ATCC 29425) were obtained from the American Type Culture Collection. *E. coli* isolates FRIK1989 and TW14588 were obtained from The Thomas S. Whittam STEC Center, Michigan State University, MI. *E. coli* isolate RM6067W was obtained from Michelle Q. Carter at the Produce Safety and Microbiology Unit, Western Regional Research Center, Agricultural Research Service, USDA. *E. coli* were grown on LB agar plates at 37°C overnight and then maintained at 4°C; fresh cultures were re-cultured every two to three weeks. Vero cells were obtained from the American Type Culture Collection (ATCC CCL-81) and cultured in DMEM (11995-065, Gibco, Thermo Fisher Scientific, Waltham, MA) supplemented with 10% FBS at 37°C and 5% CO_2_ in T75 tissue culture flasks; flasks were trypsinized and subcultured twice weekly. Bovine intestinal epithelial cells (BIECs) were obtained from Radhey Kaushik at South Dakota State University, Brookings, South Dakota. BIECs were cultured at 37°C and 5% CO_2_ in DMEM/F12 (Gibco, Thermo Fisher Scientific, Waltham, MA) supplemented with 5% FBS, 25 ng/ml mouse epidermal growth factor (PMG8041, Gibco, Thermo Fisher Scientific, Waltham, MA), 1x Insulin-Transferrin-Selenium (41400045, Gibco, Thermo Fisher Scientific, Waltham, MA), and penicillin/streptomycin (P4333, Sigma-Aldrich, St. Louis, MO); herein referred to as complete DMEM/F12 culture media. Confluent T75 flasks of BIECS were trypsinized and subcultured twice weekly.

### Growth curves of the *E. coli* isolates

Growth curves of the isolates were determined using a Bioscreen microplate reader (Growth Curves USA, Piscataway, NJ). Cultures were grown overnight in LB broth at 37°C with shaking at 200 rpm from a colony scrape off a LB agar plate. These overnight cultures were standardized to OD600 value of 0.5, and then diluted 1:100 in test media (low glucose DMEM (11054-020, Gibco, Thermo Fisher Scientific, Waltham, MA) or LB broth (L7275-500TAB, Sigma-Aldrich, St. Louis, MO)). The diluted cultures were pipetted (300 µl) into the wells of a honeycomb plate. The Bioscreen growth conditions were static growth at 37°C with wideband filter readings taken every 30 minutes for 24 hours.

### Polymorphic amplified typing sequence (PATS)

Colony lysates, prepared from each bacterial isolate cultured on LB agar plates, were each tested in triplicate to confirm the PATS profiles as described previously (21, 81–83). Briefly, this bacterial genetic fingerprinting was done using primer pairs targeting the 8 polymorphic *Xba*I-, 7 polymorphic *Avr*II-restriction enzyme sites, and the four virulence genes encoding Shiga toxin 1 and 2 (*Stx*1 and *Stx*2), Intimin-γ (*eae*), and hemolysin-A (*hly*A); hot start, touchdown PCR was used to generate amplicons from colony lysates (19, 21, 81, 83, 84). Amplicons with the putative *Avr*II-restriction enzyme sites were purified using the QIAquick PCR purification kit (Qiagen, Valencia, CA), and digested with the *Avr*II restriction enzyme (New England Biolabs, Beverly, MA) to confirm the presence of the restriction site. All reactions were analyzed by electrophoresis on 3% agarose gels stained with ethidium bromide. The presence or absence of amplicons for *Xba*I and the virulence genes was recorded using “1” and “0”, respectively. The absence of an AvrII amplicon was recorded as “0”, and the presence of restriction site with a SNP as “1”, “2” for an intact restriction site, and “3” for a restriction site duplication (19, 21, 81).

### Cattle challenge and shedding enumeration

The O157:H7 isolates were grown in 10 mL LB culture at 37°C and 190 rpm overnight, the cultures were then diluted 1:100 and grown at 37°C and 190 rpm for 7.5 hours. Next, the cultures were pelleted, resuspended in 10 mL LB supplemented with 10% glycerol, and frozen. The frozen diluted stocks (10 mL) were mixed with 90 mL phosphate-buffered saline to create inoculum for challenge. Jersey-specific dairy breed calves were purchased and were housed in climate-controlled facilities. Calves were given ad libitum access to food (alfalfa cubes hay) and water for the duration of the experiment. All animal protocols were approved by the National Animal Disease Center Animal Care and Use Committee. Eight calves per strain were orally challenged with FRIK1989 and TW14588, and seven calves per isolate were challenged with EDL933 and RM6067W. Challenge occurred orally with 10 mL of the prepared inoculum, which contained approximately 6 x 10^9^ CFU of each respective isolate. Fecal samples were collected six- or nine-days pre-challenge, on the day of challenge, and on days 1, 2, 3, 4, 5, 7, 9, 11, and 14 relative to day of challenge. Bacterial enumeration of fecal *E. coli* O157 was performed by direct and enrichment culture on selective media, as previously described (85). Briefly, non-enrichment O157 culture enumeration involved placing 10g of feces in 50ml of tryptic soy broth supplemented with cefixime (50 µg/liter), tellurite (2.5 mg/liter), and vancomycin (40 mg/liter) (TSB-CTV) with vortexing to generate a homogenous suspension. The fecal suspension was serially log diluted in sterile 0.9% saline and 100 µl spread plated on sorbitol MacConkey agar plates (SMAC) supplemented with 4-Methylumbelliferyl-β-D-glucuronide (100 mg/liter) (MUG), which were cultured overnight at 37°C. Enrichment cultivation involved culturing 10g of feces in TSB-CTV broth for 18 hours at 37°C and 150 rpm, followed by serial log dilution and spread plating as noted for the non-enrichment cultures above with the exception that SMAC-CTMV plates were used. Sorbitol negative and MUG positive colonies were subject to the *E. coli* O157 latex agglutination test kit (Thermo Fisher Scientific, Waltham, MA) to confirm serotype, with subsequent PATS typing. Recto-anal junction (RAJ) mucosal colonization by the four test *E. coli* O157 was evaluated by sampling the RAJ of individual calves with four foam tipped swabs as previously described (86). The swabs were initially placed in 10 ml of TSB-CTV media for transport back to the lab. The tube with swabs was vortexed and then an additional 40 ml of TSB-CTV media was added and mixed well. Bacterial cultivation and enumeration followed that stated above for fecal O157 *E. coli* counts. Cumulative fecal shedding and RAJ colonization was investigated using an area under the log curve analysis, and statistical differences were determined by One-Way ANOVA with Tukey’s comparison of means where *p* < 0.05 was considered significant (Rstudio 4.1.2; https://www.rstudio.com).

### DNA & library preparation

Genomic DNA (gDNA) was extracted from the culture used to inoculate calves with the DNeasy blood and tissue Genomic-tip kit (Qiagen, Hilden, Germany). gDNA quality was assessed by Nanodrop spectrophotometry (Thermo Fisher Scientific, Waltham, MA) and a Qubit fluorometer DNA broad range kit (Thermo Fisher Scientific, Waltham, MA). Nanopore libraries were prepared with rapid barcoding kit (SQK-RBK004) according to the manufacturer’s instructions (Oxford Nanopore, Oxford, U.K.). The Nextera Flex kit (Illumina, San Diego, CA) was used to prepare the genomic library for MiSeq sequencing.

### Genomic sequencing & assembly

Long-read sequencing was performed on an Oxford Nanopore MinION instrument using a FLO-MIN106 R9.4.1 flow cell for 48 hours. Bases were called and data de-multiplexed with Guppy v. 3.1.5 (87). Reads with a quality (Q) score of less than seven were removed from the analysis. Long-read sequencing data was independently assembled with the Flye assembler v2.7 (88, 89). Short-read sequencing was performed on an Illumina MiSeq instrument for 500 cycles (2 x 250) v2 kits with the Nextera Flex protocol. The resultant reads were trimmed and filtered with the BBTools software package (90). Filtered short reads were used to polish the long-read assemblies with Pilon (91), and reads were mapped to confirm the final assemblies with BBMap (90).

### *In silico* analysis of bacterial genomes

The LSPA6 lineages were assigned by a custom script (https://github.com/USDA-FSEPRU/O157LineageAssignment). Virulence genes were screened for each isolate using the Specialty Genes function of PATRIC (https://www.patricbrc.org) utilizing the Virulence Factor property and VFDB source (92, 93). Genes of interest were extracted, translated, and aligned using Geneious Prime (https://www.geneious.com/prime/) to detect polymorphisms in sequences. *In silico stx* gene subtyping was also performed in Geneious Prime utilizing published subtyping primer sequences (94).

### Phylogenetic tree

We used genomes available in the NCBI pathogen detection project (95) (version PDG000000004.3284) to place our infection genomes in the phylogenetic context of all publicly available *E. coli* O157:H7 genomes. First, all *stx-* and *eae-*positive genomes were identified and downloaded. For a genome to be considered *stx* positive, it needed to contain both the A and B subunits of either *stx1* or *stx2*, as identified by the AMRFinderPlus (96) tool (part of the NCBI pathogens pipeline). These downloaded genomes were then serotyped with the tool ‘ectyper’ (97). Only genomes with a serotype of “O157:H7” were considered for further analysis. The LSPA-6 lineage typing scheme (14, 98) was used to assign genomes into one of three lineages via a tool available at https://github.com/USDA-FSEPRU/O157LineageAssignment. To assess the large-scale phylogenetic structure of these available genomes, one genome per SNP cluster was randomly selected as a representative. In addition to these representative genomes, the reference genomes *E. coli* O157:H7 str. Sakai (GCF_000008865.2), *E. coli* O157:H7 str. 86-24 (GCF_013168095.1) and *E. coli* O157:H7 str. TW14359 (GCF_000022225.1) and the infection isolates sequenced as part of this publication were added and a pangenome was constructed using ppanggolin (99). A concatenated multiple sequence alignment was constructed from the core genome and a maximum likelihood tree was inferred using RAxML (100) using a GTRGAMMA model.

### *Ex vivo* recto-anal junction squamous epithelial (RSE) cell attachment assay

The RSE cell attachment assays were performed as previously described (53, 84, 101, 102), using bacterial isolates cultured overnight in Dulbecco’s modified Eagle’s medium with low glucose (DMEM-LG; Invitrogen, Carlsbad, CA) at 37°C without aeration, washed, and re-suspended in DMEM with no glucose (DMEM-NG; Invitrogen, Carlsbad, CA) (52, 84, 103). In addition to the test O157 isolates, *E. coli* K12 was included as a comparative control in these assays. Assays were done in triplicates, with eight technical replicates per bacterial isolate per assay. Briefly, RSE cells, collected from bovine rectal-anal junction (RAJ) tissue at necropsies, were suspended in DMEM-NG to a final concentration of 10^5^ cells/ml and mixed with bacteria at a bacteria:cell ratio of 10:1 (52, 84, 101, 102). The mixture was incubated at 37°C with aeration (110 rpm) for four hours, pelleted, washed, and reconstituted in 100 µl of double-distilled water (dH_2_O). Eight drops of the suspension (2 µl) were placed on Polysine slides (Thermo Scientific/Pierce, Rockford, IL), dried, fixed, and stained with fluorescent-tagged antibodies specific to the O157 antigen and cytokeratins within the RSE cells (52). The fluorescein isothiocyanate (FITC; green)–labelled goat anti-O157 (KPL, Gaithersburg, MD, USA) antibody targeting the O157 antigen and the mouse anti-(PAN) cytokeratins (AbD Serotec, Raleigh, NC, USA) in combination with Alexa Fluor 594 (red)–labelled goat anti-mouse IgG (H+L; F(ab')_2_ fragment) (Invitrogen) targeting the RSE cell cytokeratins were used (52). The primary rabbit anti-*E. coli* (Thermo Scientific Pierce) antibody and the secondary Alexa Fluor 488 (green) labeled goat anti-rabbit IgG (H + L; F (ab’)2 fragment) (Invitrogen) targeting the anti-*E. coli* antibody were used to detect *E. coli* K12 (52).

Attachment patterns on RSE cells were qualitatively recorded as diffuse, aggregative, or nonadherent, and quantitatively as the percentages of RSE cells with or without adhering bacteria (53, 84, 102); attachment was recorded as strongly adherent when more than 50% of RSE cells had 10 adherent bacteria, moderately adherent when 50% or less of the RSE cells had 1 to 10 adherent bacteria, and nonadherent when less than 50% of the RSE cells had only 1 to 5 adherent bacteria. RSE cells with no added bacteria were subjected to the assay procedure and used as negative controls to confirm absence of pre-existing O157 bacteria. Quantitative data was compared between isolates for statistical significance using one-way ANOVA with Tukey’s multiple comparisons test; *p* < 0.05 was considered significant (GraphPad Prism version 8.0.0, GraphPad Software, San Diego, CA).

### *In vitro* cell attachment assay

*In vitro* cell attachment phenotypes were determined using bovine intestinal epithelial cells (BIECs). Confluent BIECs from a three-day T75 flask were trypsinized and diluted to 100,000 cells/ml in complete DMEM/F12 culture media and 200 µl of this suspension was pipetted into the wells of a sterile 96 well tissue culture plate (20,000 cells/well). These tissue culture plates were then cultivated at 37°C and 5% CO_2_ for two days to obtain confluent monolayers. A colony scrape taken from an LB agar plate of *E. coli* was inoculated into five ml of low glucose DMEM (11054-020, Gibco, Thermo Fisher Scientific, Waltham, MA) and grown overnight at 37°C with shaking at 200 rpm. Following overnight growth, the OD600 of these cultures was measured and adjusted to an OD600 1.0 and a 1:10 dilution in low glucose DMEM was made. The media on the BIECs was removed and replaced with 200 µl of low glucose DMEM, then 10 µl of the diluted *E. coli* suspensions were added to the BIECs, and the tissue culture plates were then centrifuged at 250 rcf for 10 minutes at room temperature to pellet the bacteria onto the BIECs. The tissue cultures plates were then placed in an incubator at 37°C and 5% CO_2_ for three hours to allow for the *E. coli* to attach to the BIECs. Following the three-hour co-incubation, the media with non-adherent bacteria was removed and the wells were washed four times with PBS to further remove non-adherent bacteria. Then 200 µl of Triton X-100 (1.0% in PBS) was added for 10 minutes to lyse the BIECs and release the adherent bacteria; a serial log dilution was made of these in PBS. The serial dilutions were spotted (3 x 20 µl spots per dilution) onto LB agar plates and grown overnight at 37°C to determine adherent CFU counts. Statistical analysis was done by One-Way ANOVA with Tukey’s comparison of means to identify significance in adherent phenotype between *E. coli* isolates (GraphPad Prism version 9.0.1, GraphPad Software, San Diego, CA).

### Biofilm production

Biofilm production was determined through crystal violet staining and the viability reagent resazurin on polystyrene microtiter plates; crystal violet was employed to assess biofilm biomass, while resazurin was used to give an indication of cellular viability within the biofilms. Overnight *E. coli* cultures grown in YESCA broth (0.1% yeast extract and 1% casamino acids) from colony scrapes were diluted to identical OD600 0.5 suspensions and diluted 1:10 in test media (YESCA or Brain Heart Infusion (BHI, CM1135, Oxoid, Basingstoke, UK)). Two hundred microliters of the suspensions in test media were pipetted into sterile 96 well high binding flat bottom microtiter plates and maintained at either 26°C or 37°C for five days. To quantify biofilm production by crystal violet staining, the media with non-biofilm embedded bacteria was removed and the wells washed with PBS two times. The biofilms in the microtiter plates were then heat fixed at 80°C for 30 minutes. After the plates had cooled, 200 µl of crystal violet (0.1% in H_2_O) was added for 30 minutes at room temperature. The stain was removed, and the wells washed four times with PBS to remove excess stain. After the final PBS wash was removed, the plates were left at room temperature overnight to allow the wells to completely dry. The bound stain was solubilized by adding 200 µl 95% ethanol and absorbance was measured at 590 nm with a BioTek Synergy HTX multimode reader (Agilent, Santa Clara, CA). To quantify biofilm production using resazurin, the growth media was removed, and the wells of the microtiter plate were washed two times with PBS to remove non-embedded bacteria. Two hundred microliters of HBSS were then added to the wells along with 20 µl of 0.3 mg/ml resazurin (R-2127, Sigma, St. Louis, MO). The plates were held at 37°C for 45 minutes to allow for the reduction of the resazurin, at which time the resazurin reduction was measured at 530 excitation and 590 emission with a Biotek Synergy HTX multimode reader (Agilent, Santa Clara, CA).

### Congo Red binding of the *E. coli* isolates

Twenty microliter drops of a 1:10 dilution of OD600 0.5 bacterial suspensions in PBS were spotted onto Congo red agar (YESCA agar (0.1% yeast extract, 1% casamino acids, 15g agar/L), 40 µg/ml Congo red dye, and 6.24 µg/ml Coomassie brilliant blue G-250) and cultivated at 26°C for 48 hours (24). Representative photographs were taken; red colonies indicate Congo red binding to curli fimbria, which is associated with adhesion and biofilm formation.

### Toxin production

Shiga toxin production by O157 isolates was determined through the cytotoxic effects of bacterial supernatants on Vero cell (33). Resazurin was used to measure the viability of the treated Vero cells. Vero cells were maintained in DMEM (11995-065, Gibco, Thermo Fisher Scientific, Waltham, MA) supplemented with 10% FBS unless otherwise noted. Two hundred microliters of Vero cells at a cell density of 1.0 x 10^5^ cells/ml were seeded into sterile 96 well tissue culture plates to achieve 20,000 Vero cells per well. These culture plates with Vero cells were cultivated at 37°C and 5% CO_2_ for two days to allow for confluent monolayers to develop. *E. coli* was grown overnight at 37°C and 200 rpm in LB broth (L7275-500TAB, Sigma-Aldrich, St. Louis, MO) from a colony scrape taken off an LB agar plate. The overnight cultures were adjusted to the same OD600 0.5 absorbance and diluted 1:100 in low glucose DMEM (11054-020, Gibco, Thermo Fisher Scientific, Waltham, MA); these cultures were incubated statically for six hours at 37°C. Six hours was chosen for toxin induction as this time period has previously been shown to produce high levels of Shiga toxins (104, 105). These cultures were then centrifuged at 1600 rcf at room temperature for 10 minutes. The supernatants were collected and passed through sterile 0.2 µm syringe filters. These filtered supernatants were then tested for cytotoxicity on Vero cells. A dilution series of the supernatants was made in DMEM supplemented with 10% FBS. The media from the confluent Vero cell culture plates was removed and replaced with 100 µl DMEM supplemented with 10% FBS, followed by the addition of 100 µl from the bacterial supernatant dilution series; final dilutions tested were 1:4, 1:40, 1:400, and 1:4000. Controls included Vero cells treated with dilution series of low glucose DMEM sans bacteria cultivation. The Vero cells were then maintained at 37°C and 5% CO_2_ for 48 hours, at which time the media from the Vero cells was removed and replaced with 200 µl phenol red free DMEM supplemented with 5% FBS; reduced serum was used as excessive serum protein has been shown to interfere with resazurin measurements (106). Twenty microliters of 0.3 mg/ml resazurin (R-2127, Sigma, St. Louis, MO) was then added and the culture plates were incubated for 150 minutes at 37°C and 5% CO_2_ to allow for reduction of the resazurin. The reduction of resazurin by viable Vero cells was measured with 530 emission and 590 excitation settings on a BioTek Synergy HTX multimode reader (Agilent, Santa Clara, CA) with Gen5 version 3.10 acquisition software. Using the background subtracted fluorescent readings the viability of the treatment groups was made relative to the controls ((Test/Control)*100). Statistical analysis was performed by One-Way ANOVA with Tukey’s comparison of means to identify significance in toxin production phenotype between *E. coli* isolates (GraphPad Prism version 9.0.1, GraphPad Software, San Diego, CA).

As stx2 is associated with worse clinical outcome, anti-stx1 antibody was employed to neutralize the cytotoxic activity of stx1 produced by these O157 isolates in these Vero cell cytotoxicity assays. Bacterial supernatants for testing were produced as stated in the preceding paragraph. Monoclonal anti-stx1 antibody (13C4, Invitrogen, Thermo Fisher Scientific, Waltham, MA) was incubated with 1:20 dilutions of the bacterial supernatants in cell culture media at an antibody ratio of 1:50 for two hours at 37°C with shaking at 100 rpm. The remaining produce followed that outlined in the previous paragraph, with the inclusion of a set of control Vero cells that received the monoclonal antibodies in cell culture media sans bacterial supernatants.

## Data availability

Sequencing data associated with the infection isolates’ genome are available through NCBI bioproject accession PRJNA884395.

## Acknowledgments

We would like to thank Bryan Wheeler, Zahra F. Bond, and Lindsey Andersen for their assistance with this research. This research was supported by appropriated funds from USDA-ARS CRIS 5030-32000-225-00D and an appointment to the Agricultural Research Service (ARS) Research Participation Program administered by the Oak Ridge Institute for Science and Education (ORISE) through an interagency agreement between the United States Department of Energy (DOE) and the United States Department of Agriculture (USDA). ORISE is managed by Oak Ridge Associated Universities (ORAU) under DOE contract number DE-SC0014664. This research used resources provided by the SCINet project of the USDA Agricultural Research Service, ARS project number 0500-00093-001-00-D. All opinions expressed in this paper are the authors’ and do not necessarily reflect the policies and views of USDA, ARS, DOE, or ORAU/ORISE.

---

**FIG S1.**
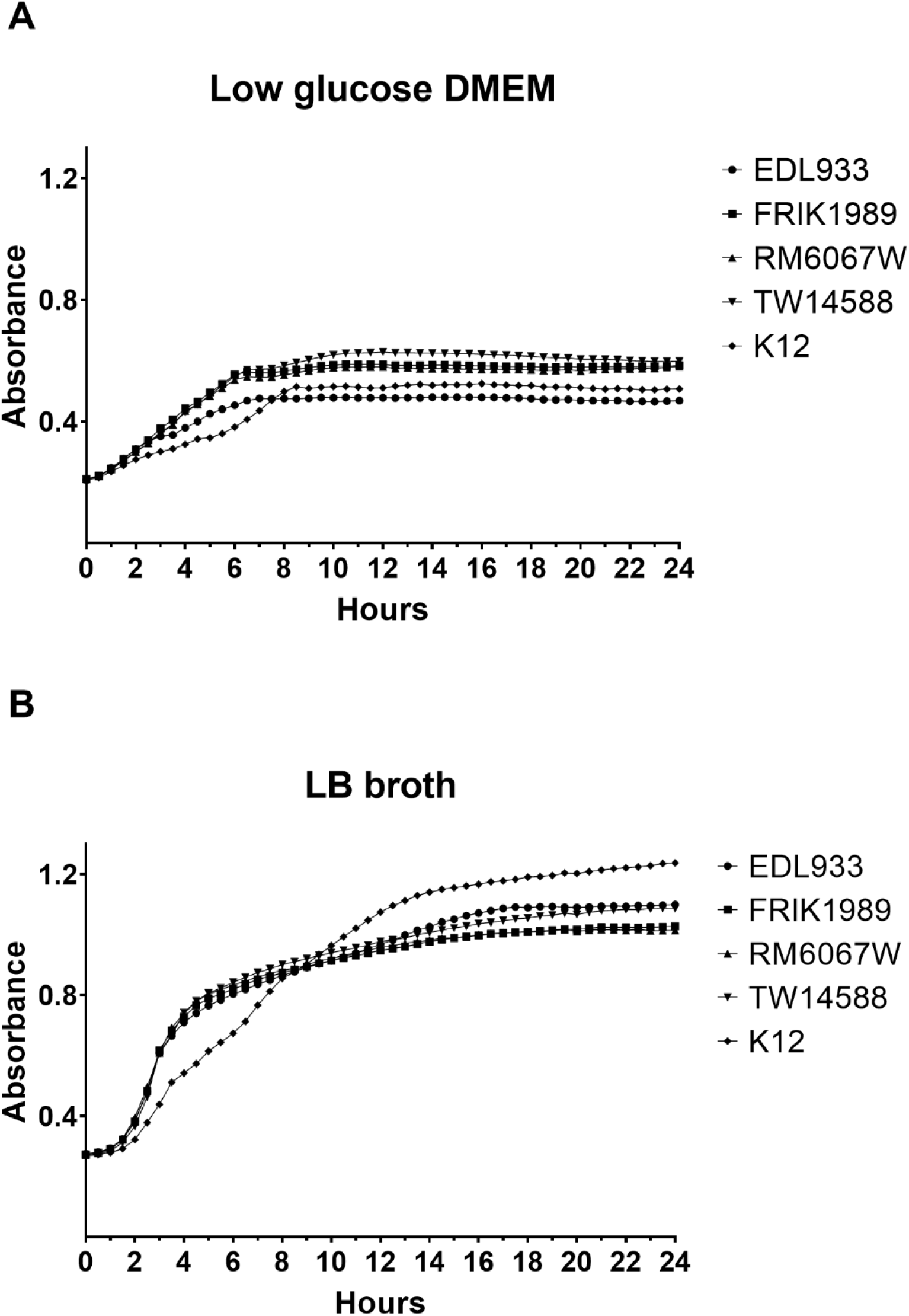
Growth profiles of *E. coli* O157 isolates in low glucose DMEM and LB broth. Absorbance readings were measured on a Bioscreen microplate reader for *E. coli* isolates EDL933, FRIK1989, RM6067W, TW14588, and K12 grown statically at 37°C. Measurements were taken every 30 minutes using the wideband filter for 24 hours. **(A)**. Growth curve in low glucose DMEM. **(B)**. Growth curve in LB broth.

**FIG S2.**
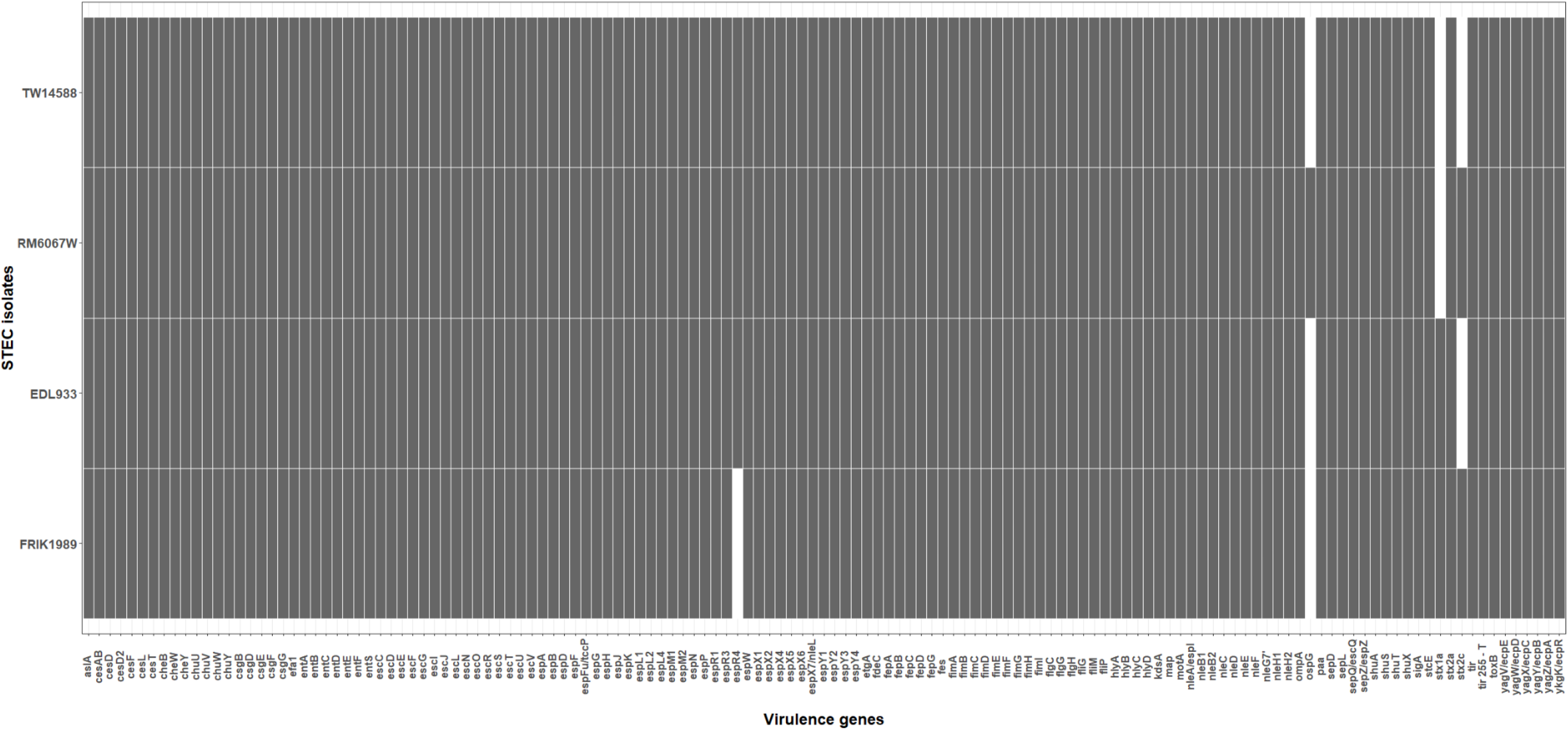
Heat map depicting major virulence genes/SNPs shared by these isolates. Virulence genes were annotated for these isolates’ genomes through PATRIC 3.6.12 (https://www.patricbrc.org) utilizing the Virulence Factor property and VFBD source within the Specialty Genes function. Further *in silico* analysis using Geneious Prime (https://www.geneious.com/prime/) was employed to ascertain *tir* 255 T/A allele and subtype *stx* genes. Grey indicates presence and white indicate absence of virulence trait.

**FIG S3.**
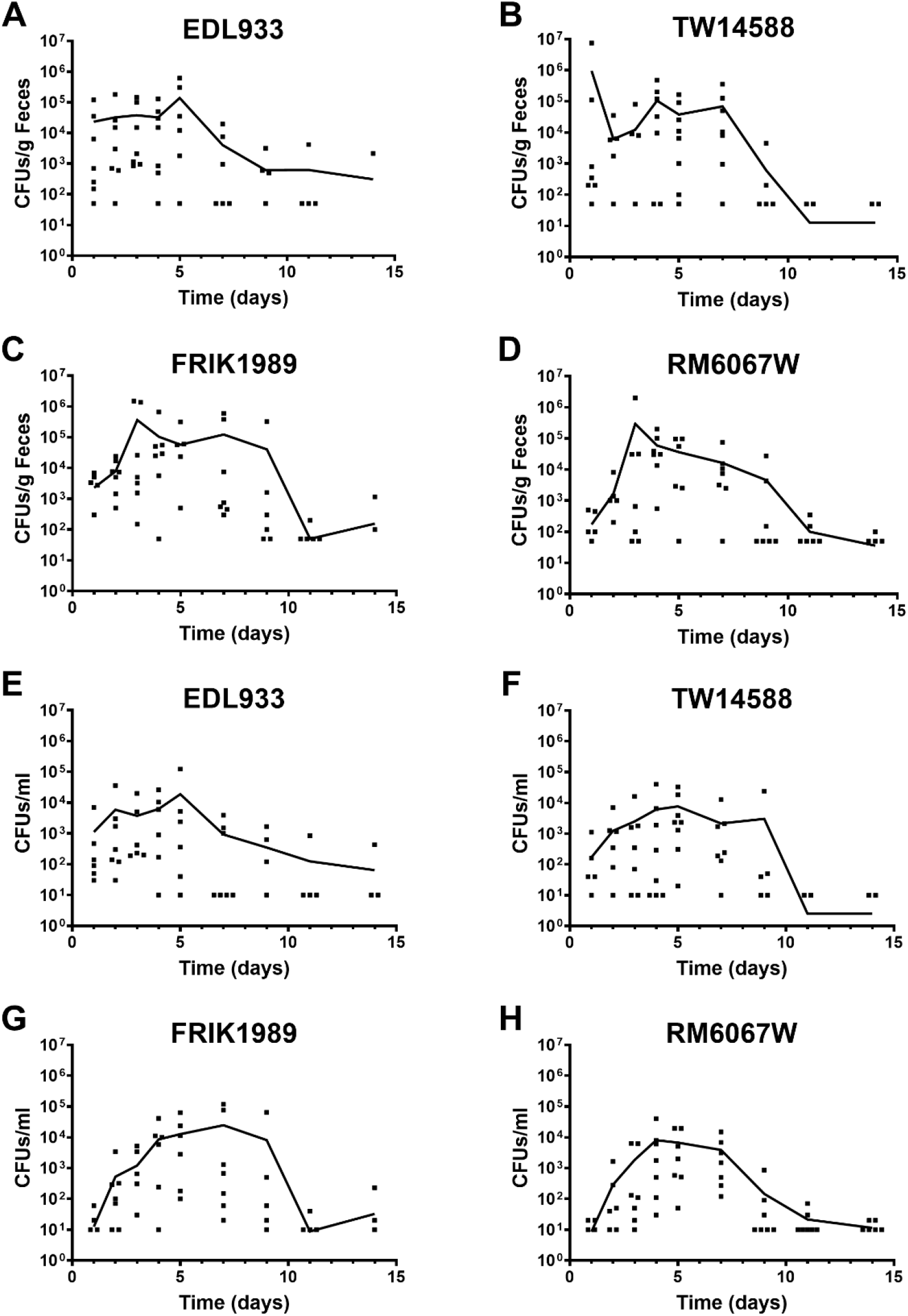
Time course of *E. coli* O157 fecal shedding and RAMS colonization. Jersey calves were orally inoculated with 6 x 10^9^ CFUs of one of the listed *E. coli* O157 isolates (EDL933, TW14588, FRIK1989, or RM6067W). Feces and rectal anal junction mucosa swabs were collected over the course of 14 days to enumerate fecal shedding and colonization at the rectal anal junction by these isolates. Polymorphic Amplified Typing Sequences (PATS) was used on *E. coli* O157 isolates collected to verify that those recovered were from the inoculated isolates and not commensals. Samples were taken from 3 or 4 animals per timepoint for each *E. coli* O157 isolate cohort over the course of two experiments. The plotted line represents the mean of the individual data points from each day; individual data points in which no CFUs were recovered are not plotted due to a log axis. STEC isolate is noted above its corresponding graph. **(A-D).** Fecal shedding. **(E-H).** Rectal anal junction mucosa swabs.

**FIG S4.**
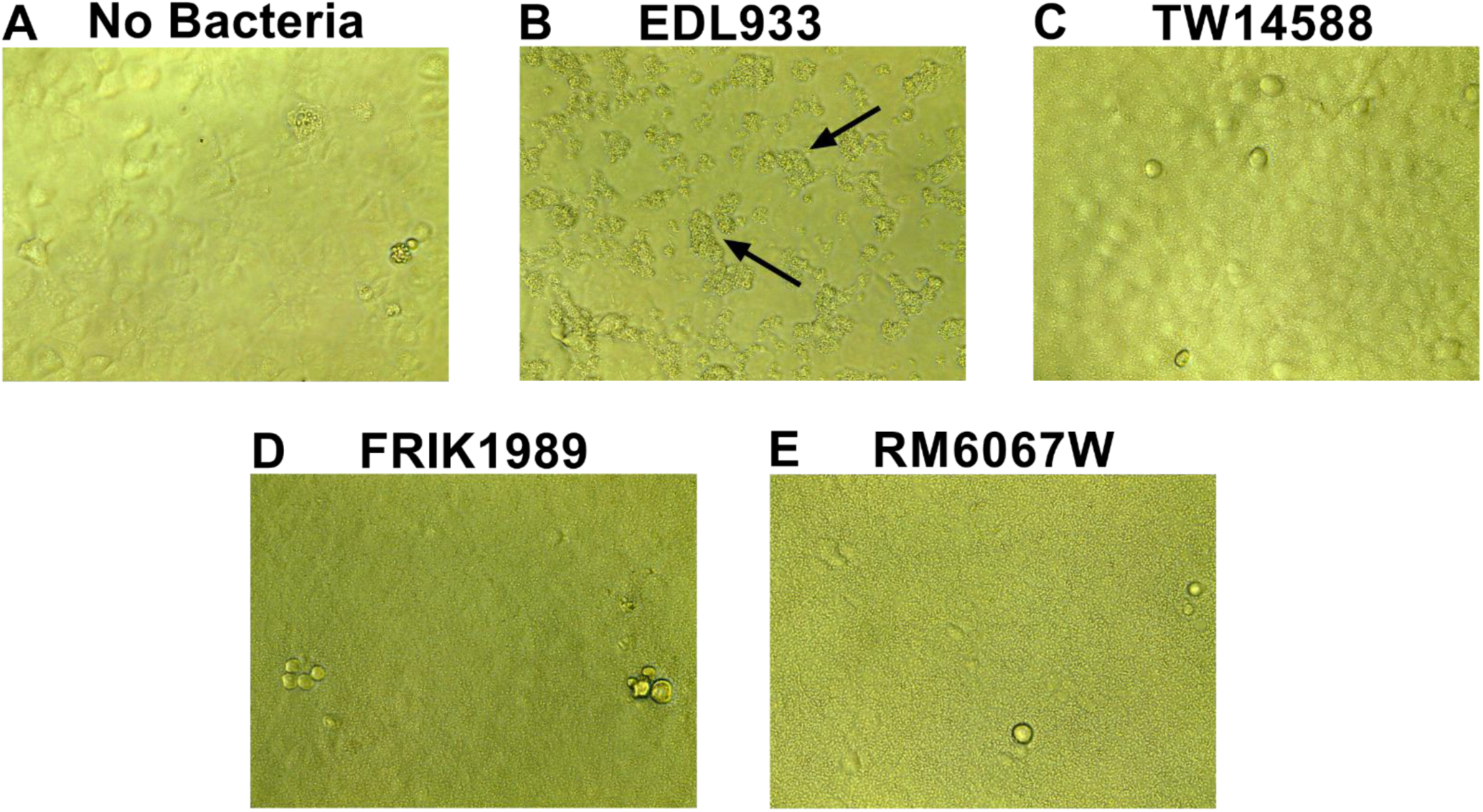
Representative brightfield microscope images displaying altered attachment phenotype of *E. coli* O157 isolate EDL933 on bovine intestinal epithelial cells (BIEC). *E. coli* was added to BIEC cells and incubated for 3 h to assess cell attachment for each respective isolate. Images were captured prior to any washing steps that would have removed nonadherent bacteria. EDL933 formed clustered microcolonies on the BIECs (black arrows) which is absent from the other *E. coli* isolates tested. **(A).** No bacteria control. **(B).** EDL933. **(C)**. TW14588. **(D).** FRIK1989. **(E).** RM6067W.

**FIG S5.**
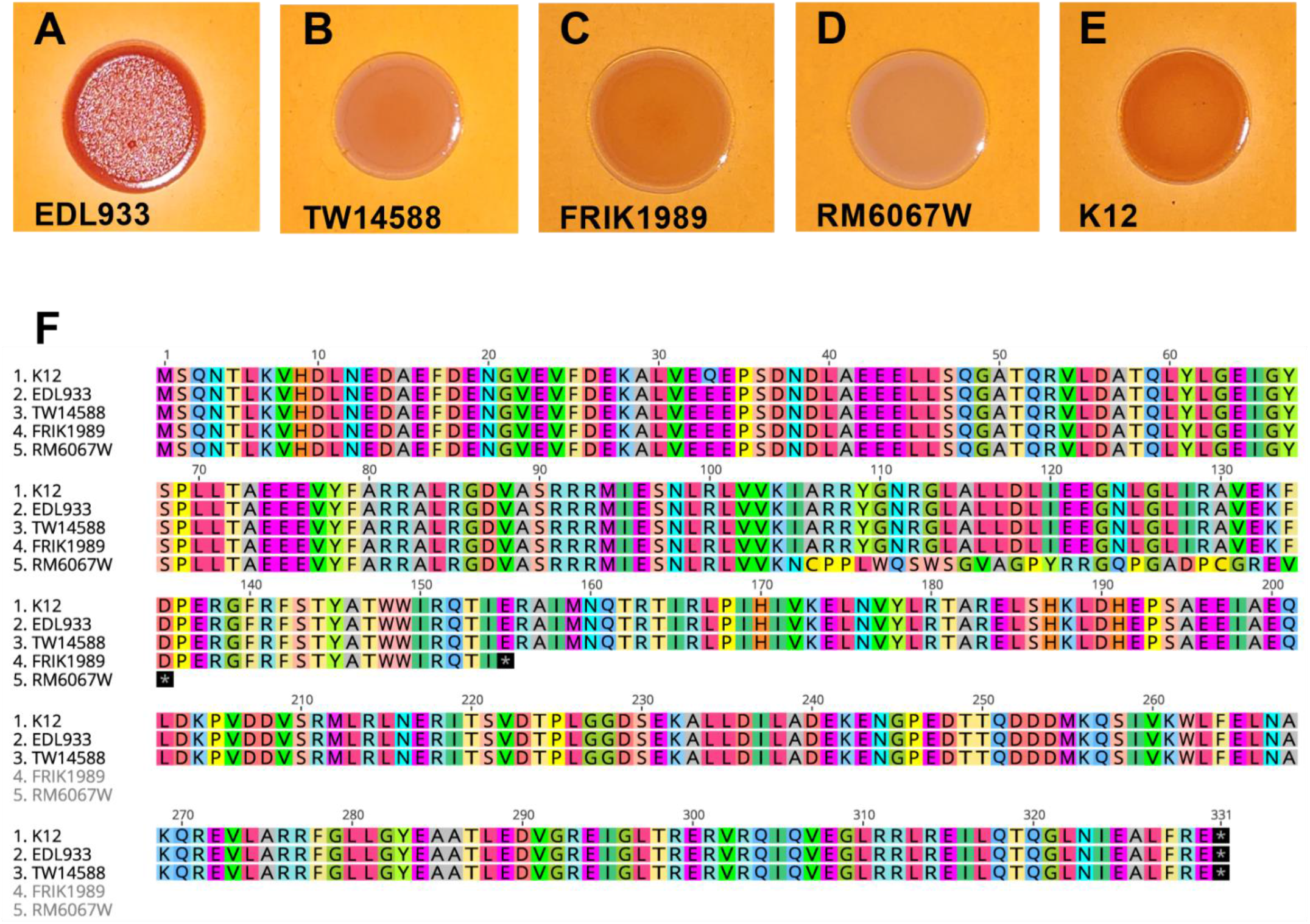
(A-E). *E. Coli* O157 phenotype on Congo Red Agar and relationship to RpoS protein sequence. A 0.02 mL drop of a 10^-1 dilution of an OD_600_ 0.5 *E. coli* suspension was spotted onto Congo Red agar and incubated for 2 days at 26 °C. **(A).** EDL933. **(B).** TW14588. **(C)**. FRIK1989. **(D).** RM6067W. **(E).** K12. Red colonies indicate curli fimbria expression. **(F).** Sequence alignment of the regulatory protein RpoS for indicated *E. coli* isolates. RpoS amino acid sequences were aligned and translated using Geneious Prime (https://www.geneious.com/prime/); an * indicates the presence of a stop codon. The *rpoS* gene sequence used to align these genes in Geneious Prime was derived from the GenBank annotated *rpoS* gene for *E. coli* MG1655 accession number NP_417221.

**Table S1.**
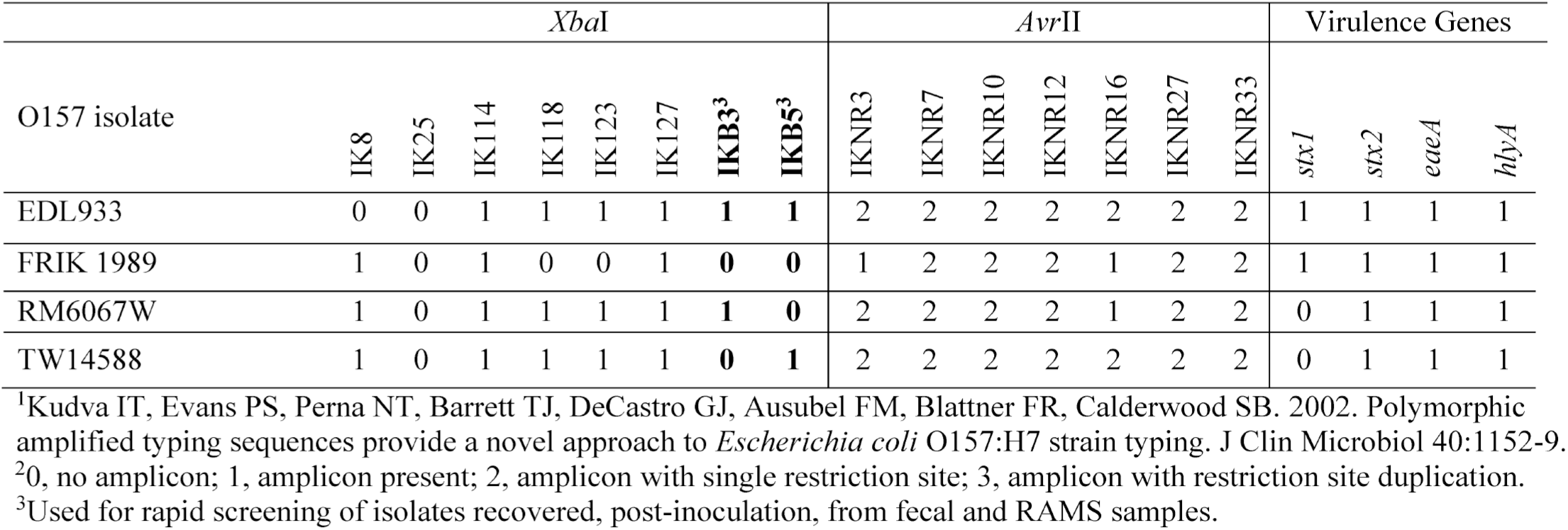
Polymorphic Amplified Typing Sequence (PATS) profiles^1^,^2^ ofO157 isolates used in this study.

